# GPR183 mediates the capacity of the novel CD47-CD19 bispecific antibody TG-1801 to heighten ublituximab-umbralisib (U2) anti-lymphoma activity

**DOI:** 10.1101/2022.03.31.486558

**Authors:** Marcelo Lima Ribeiro, Núria Profitós-Pelejà, Juliana Carvalho Santos, Pedro Blecua, Diana Reyes Garau, Marc Armengol, Miranda Fernández-Serrano, Hari P. Miskin, Francesc Bosch, Manel Esteller, Emmanuel Normant, Gael Roué

**Affiliations:** Lymphoma Translational Group, Josep Carreras Leukemia Research Institute, Badalona, Spain; Cancer Epigenetics Group, Josep Carreras Leukemia Research Institute, Badalona, Spain; Laboratory of Immunopharmacology and Molecular Biology, Sao Francisco University Medical School, Braganca Paulista, São Paulo, Brazil; Autonomous University of Barcelona, Barcelona, Spain; TG Therapeutics, New York, NY, USA; Department of Hematology, Vall d’Hebron University Hospital, Barcelona, Spain; Experimental Hematology, Vall d’Hebron Institute of Oncology, Barcelona, Spain; Centro de Investigación Biomédica en Red de Cáncer (CIBERONC), Instituto de Salud Carlos III, Barcelona, Spain; Instituciò Catalana de Recerca i Estudis Avançats (ICREA), Barcelona, Spain

## Abstract

Targeted therapies have considerably improved the survival rate of B-cell non-Hodgkin lymphoma (B-NHL) patients in the last decade; however, most subtypes remain incurable. TG-1801, a bispecific antibody that targets CD47 selectively on CD19+ B-cells, is under clinical evaluation in relapsed/refractory B-NHL patients either as a single-agent or in combination with ublituximab, a CD20 antibody, which is also being combined with the PI3Kδ/CK1e inhibitor, umbralisib (“U2”-regimen). In this study, we demonstrated that TG-1801 potentiates ublituximab-mediated antibody-dependent cell death (ADCC) and antibody-dependent cell phagocytosis (ADCP), leading to an additive anti-tumour effect of the TG-1801/U2 combination in B-NHL co-cultures. Accordingly, in a B-NHL xenotransplant model, the triplet achieved a 93% tumour growth inhibition, with 40% of the animals remaining tumour-free 35 days after the last dosing. Transcriptomic analysis further uncovered the upregulation of the G protein-coupled receptor, GPR183, as a crucial event associated with TG-1801/U2 synergism, while pharmacological blockade or genetic depletion of this factor impaired ADCP initiation, as well as cytoskeleton remodelling and cell migration, in B-NHL cultures exposed to the drug combination. Thus, our results set the preclinical rationale and support a combination strategy of TG-1801 with PI3Kδ- and CD20-targeting agents in patients with B-NHL.

## Introduction

Cluster of differentiation 47 (CD47), also known as integrin-associated protein (IAP), is a cell surface receptor that is part of the immunoglobulin superfamily and interacts with the macrophage receptor signal regulatory protein-alpha (SIRPα). This interaction sends a “do-not-eat-me” signal to macrophages, which mediates immune evasion in several types of cancers (Barclay and Van den Berg, 2014). High levels of CD47 have been observed in both lymphoid and myeloid neoplasm in which this factor is both an adverse prognostic indicator and a valid anti-cancer target with several therapeutic antibodies currently being tested in clinical trials. In B-cell lymphoma, these trials frequently involve a combination with an anti-CD20 therapy, to ensure a proper engagement of the Fc receptors at the surface of macrophages and natural killer (NK) effector cells. The anti-CD20 mAb rituximab has been the most common IgG1 antibody tested in this setting, and has demonstrated combinatorial activity in both indolent and aggressive B-cell lymphomas (Armengol et al., 2021; Matlung et al., 2017). However, as CD47 is widely expressed on the surface of a broad range of cell types, including erythrocytes and platelets, a major limitation of CD47 blocking agents is the target-mediated drug disposition and the potential side effects, which include anaemia or thrombocytopenia.

TG-1801 is a novel bi-specific antibody that binds to CD47 with sub-micromolar affinity and to CD19 with a sub-nanomolar affinity. This thousand-fold difference between its affinity to CD19 and CD47 allows TG-1801 to bind selectively to CD19-positive B cells but not CD19-negative red blood cells or platelets (Buatois et al., 2018; Hatterer et al., 2020). TG-1801 is currently being tested clinically as a single agent and in combination with the glyco-engineered CD20 antibody ublituximab in patients with relapsed/refractory B-cell lymphoma. Ublituximab, in association with the dual PI3Kδ/CK1ε inhibitor umbralisib (a combination named U2), has demonstrated clinical activity in B-cell lymphoma, and a biologic license application for U2 in chronic lymphocytic leukaemia (CLL) has been recently accepted for review by the FDA (Lunning et al., 2019). Here we studied the triplet combination of TG-1801 and U2 in *in vitro* and *in vivo* models of aggressive B-cell lymphoma.

## Results and Discussion

### CD47 blockade in CD19+ cells potentiates the anti-tumour effect of U2 regimen

To determine the working concentrations of TG-1801 *in vitro*, we first developed a CD47 occupancy assay using the Burkitt lymphoma (BL) cell line Raji. In this assay, TG-1801 reached a plateau of 48% CD47 occupancy at a dose as low as 20 ng/mL, slightly lower than the 67% occupancy achieved by the first-in-class CD47 blocking mAb, B6H12 (Mateo et al., 1999) (Figure 1A). This difference may potentially be explained by the lower level of expression of CD19 compared to CD47 in the BL cell line (Figure 1 - figure supplement 1). As expected, TG-1801, but not B6H12-mediated target occupancy, was highly dependent on CD19 expression, as shown in the T-ALL-derived, CD19 negative, Jurkat cells, where no significant CD47 occupancy was detected with TG-1801 at doses as high as 2 μg/mL, contrasting with the sustained binding of B6H12 (Figure 1A). A panel of human B-cell lymphoma cell lines (n=10) with different levels of CD47 and CD19 expression, were cultured in the presence of M1-polarized primary macrophages or primary circulating PBMCs from healthy donors as a source of effector cells, to assess the TG-1801 antibody-dependent cell phagocytosis (ADCP) or antibody-dependent cell cytotoxicity (ADCC), respectively. As shown in Figure 1 - figure supplement 2, ADCP and ADCC were both increased 2-6 fold after CD47 ligation in these cells, and both appeared to be related to neither the expression levels of CD47 nor the CD47/CD19 ratios, in accordance with previous reports (Buatois et al., 2018). Interestingly, the analysis of the pro-cytotoxic or pro-phagocytic activities of the triplet regimen in five representative B-NHL cell lines, including two DLBCL (Pfeiffer and Karpas-422), two BL (Raji and Daudi), and one FL (RL), showed an improvement in these activities when compared to the TG-1801 single-agent treatment (Figure 1B). In line with this observation, in Raji tumour-bearing mice, all molecules were active as single agents (88%, 76% and 50% tumour growth inhibition (TGI) with ublituximab, TG-1801 or umbralisib, respectively, Figure 1 - figure supplement 3), and the activity of the triplet was higher (93% TGI) after a 17-day treatment. Furthermore, 40% of the mice remained tumour-free 35 days after the last dose in the triplet arm with no detectable toxicity (Figures 1C and 1D, and data not shown). These *in vitro* and *in vivo* data suggested that the addition of TG-1801 to the U2 combination induced a mechanism that promoted a stronger innate immune response.

**Figure 1.**
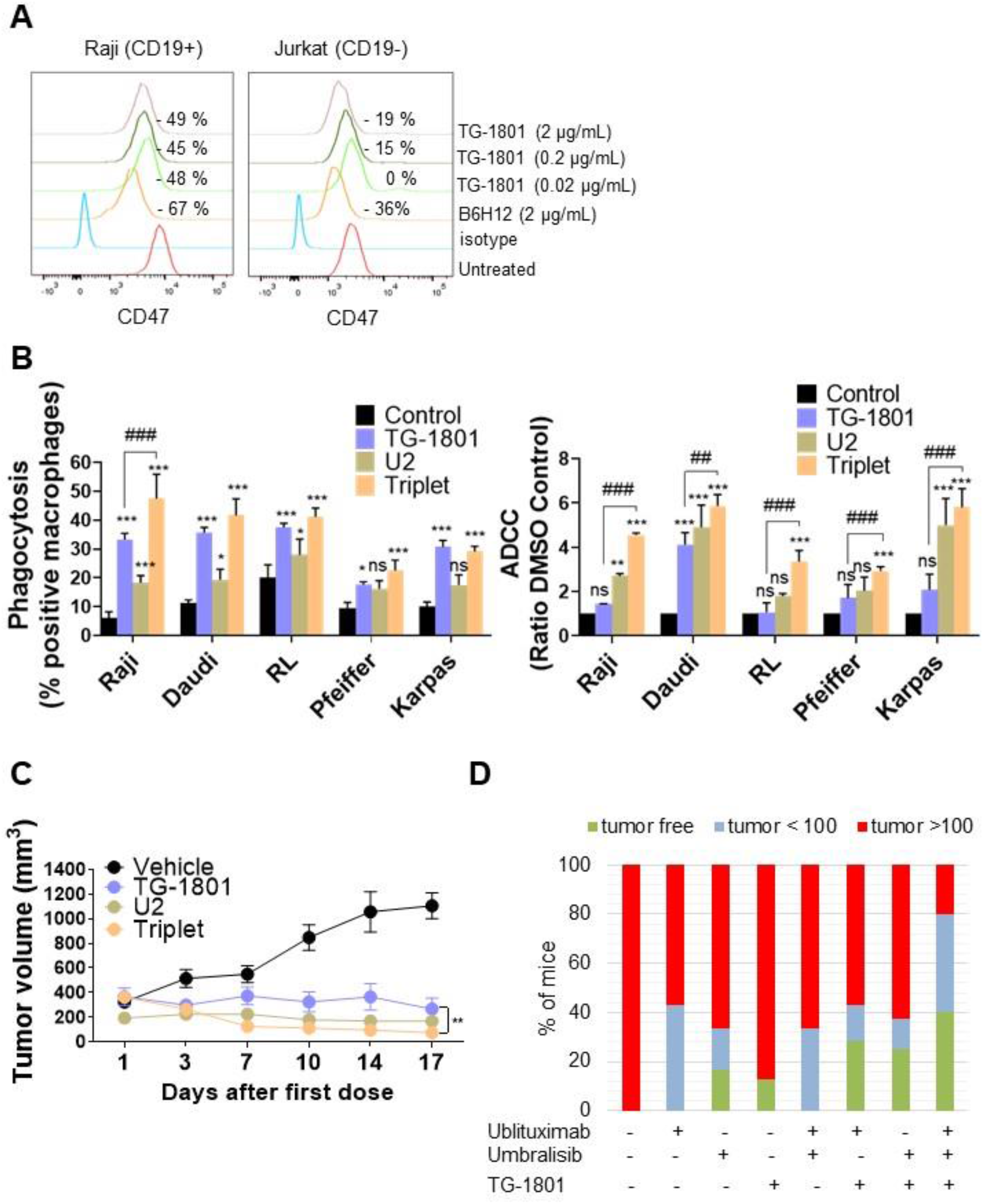
U2 regimen cooperates with CD47 blockade in *in vitro* and *in vivo* models of B-cell lymphoma. A) FACS-mediated CD47 occupancy assay in the CD19+ Burkitt lymphoma cell line Raji, and in the CD19-T-ALL cells Jurkat. Results showed a CD19-dependent, optimal competition of 20 ng/mL TG-1801 with the PE-labelled anti-CD47 antibody (N=3). B) ADCP (left panel) and ADCC (right panel) activities were assessed in five representative B-cell lymphoma cell lines (N=3). Values are expressed as mean ± SD. C) TG-1801, U2 and the triplet combination were dosed orally in the Raji xenograft model. D) Mice with no tumour or low tumour size were kept alive for another 35 days. At day 52 all the mice either tumour-free (green) or bearing a small tumour (blue, < 100 mm^3^) were alive. The TG-1801-U2 combo group showed an increased number of tumour free or low tumour burden bearing mice (green and blue bars). * *p*<0.05, ** *p*<0.01, *** *p*<0.001, when compared to control group. (N= 8-6 mice per group) ^###^ *p*<0.001 when compared to TG-1801 alone. ns = non-significant.

### GPR183 is upregulated in response to TG-1801/U2 combination treatment

To uncover the mechanisms underlying the superior effect of the triplet, transcriptomic analyses were carried out on a set of six samples that included Raji xenograft tumours (n=2) and CD20+ cells isolated from Raji, Daudi and two BL primary samples co-cultured with the bone marrow-derived stromal cell line (BMSC) stromaNKtert (Dlouhy et al., 2020), M2-polarized primary macrophages, primary circulating PBMCs, and either TG-1801 or the TG-1801+U2 triplet. As shown in Figure 2A, a total of 20 genes were significantly up- or down-regulated in the triplet compared to TG-1801 treatment in all six samples. A Gene Set Enrichment Analysis (GSEA) identified inflammatory (NES = 2.43, FDR = 0) and TNFα-driven signatures (NES = 2.43, FDR = 0) as predominantly activated upon treatment with the triple combination, when compared to TG-1801 single agent therapy, suggesting that the stronger activity of the triplet was based on the activation of an immune-related anti-tumour effect. In the heatmap showing a set of genes strongly activated in all six samples after the triplet treatment (Figure 2B), the highest up-regulated gene in both signatures was the G protein-coupled receptor 183 (GPR183, also known as Epstein-Barr virus (EBV)-induced G protein-coupled receptor 2, EBI2). GPR183 was also one of the 20 upregulated genes previously identified in the volcano plot (Figure 2A). An increase in GPR183 expression after U2 addition to TG-1801 was confirmed *in vitro* by qPCR analysis in the four cell lines-derived samples (Figures 2C and 2D), by western blot in the Raji and Daudi cell lines (Figure 2E and source data 1), and by immunohistochemistry (IHC) in tumour specimen from the Raji xenograft model (Figure 2F). GPR183 was first identified by sequence similarity as a GPCR induced by EBV infection. This pro-inflammatory receptor, which upregulation is associated with a better prognosis of DLBCL patients treated with the standard immunochemotherapeutic (R-CHOP) regimen according to published gene array database (gse10846; R2: Genomics Analysis and Visualization Platform (http://r2.amc.nl http://r2platform.com)), plays an important role in B-cell motility and positioning during the germinal centre reaction (Liu et al., 2011; Pereira et al., 2009), and the gradient of its natural ligand, oxysterol, acts as a chemoattractant of GPR183+ cells. Accordingly, among a panel of eight genes identified besides GPR183 in the two inflammation gene signatures, the transcript of the sole chemoattractant gene *CCL20* was upregulated by the triple combination in the three *in vitro* and *in vivo* models (Figures 2C and 2D). Also, increased tumour infiltration of mouse macrophages (F4/80 IHC staining) together with a reduction in mitotic index (histone H3-pSer10 IHC staining) were observed in triplet-treated tumours when compared to either TG-1801 or U2 arms (Figure 2F).

**Figure 2.**
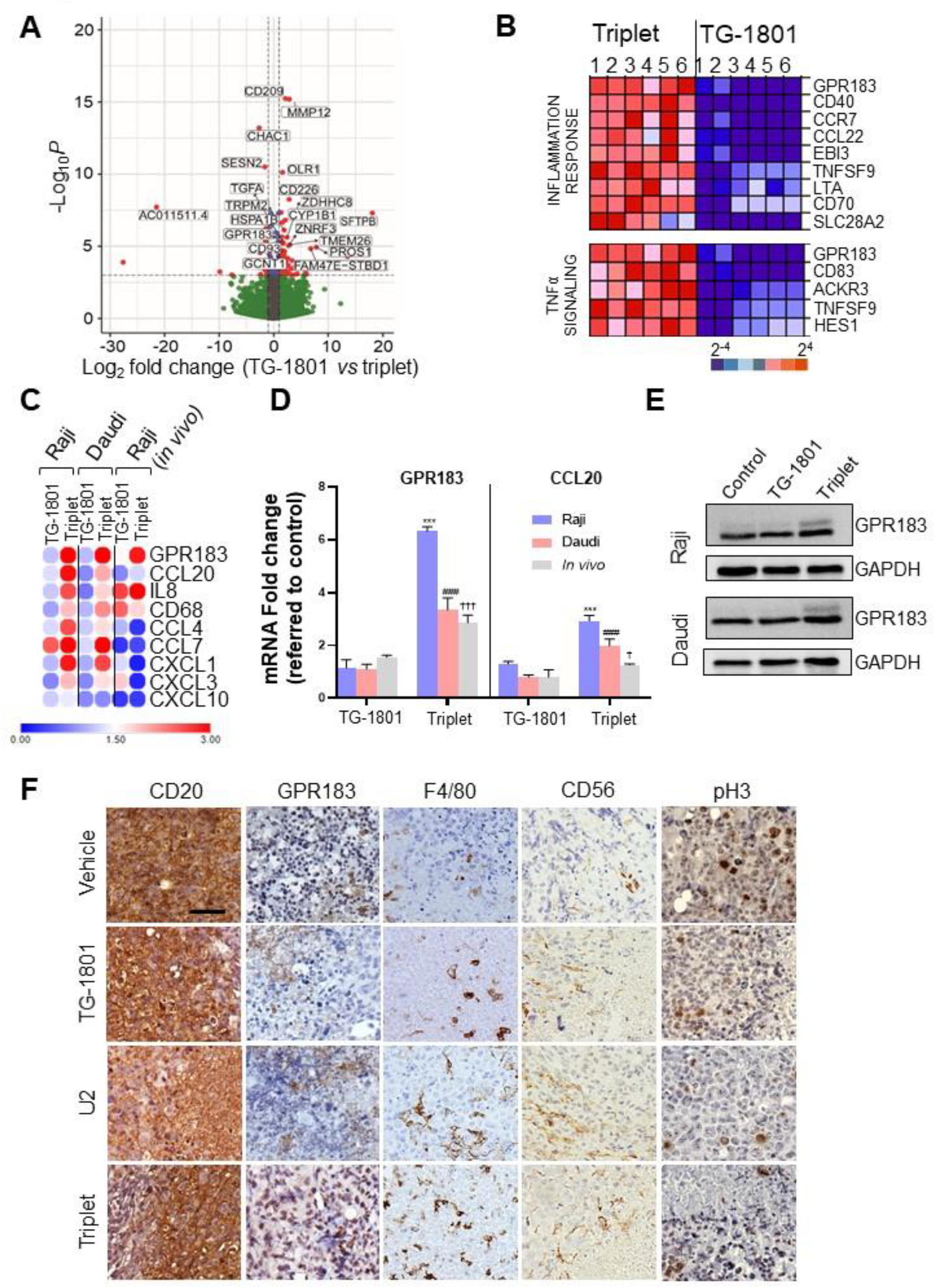
Upregulation of GPR183 is a hallmark of the response to U2 and TG-1801 combination *in vitro* and *in vivo*. A) Volcano plot showing the most relevant significantly differentially expressed genes between triplet and TG-1801 treatments. Genes that underwent the same modulation in the three *in vitro* and *in vivo* models (N=20) have been labelled. B) Gene set enrichment analysis was performed using the GSEA software to analyse the enriched gene sets in the triplet samples compared to the samples treated with TG-1801. Samples were sorted from left to right: 1-Raji, 2-Daudi, 3-4-BL primary samples and 5-6 CD20+ cells isolated from 2 representative Raji xenograft specimens. C) GPR183 upregulation was confirmed in these samples and compared to the 11 inflammatory genes extracted from GSEA analysis. Data are presented in fold-change related to the control (N=3). Clustering was performed using Morpheus (hierarchical, one minus Pearson correlation) available at https://software.broadinstitute.org/morpheus/. D) *GPR183* and *CCL20* transcript levels followed the same evolution *in vitro* and *in vivo* according to the different treatment regimens. E) Immunoblot evaluation of GPR183 (SantaCruz, #sc-514342) in both Raji and Daudi showed an increased GPR183 protein expression after the combination treatment (N=3). F) Immunohistochemistry (IHC) labelling of CD20 (Clone L26, Sigma-Aldrich), GPR183 (Clone G-12, Santa Cruz), F4/80 (Clone SP115, Abcam), Histone H3-pSer10 (Clone E173, Abcam) and CD56/NCAM-1 (Clone EPR1827, Abcam) in tissue sections from tumour specimens (N=3). *** *p*<0.001, ^###^ *p*<0.001, and ^†^ *p*<0.05 and ^†††^ *p*<0.001 when compared to control group in Raji (*in vitro*), Daudi (*in vitro*) and Raji (*in vivo*) models, respectively. ns = non-significant. Figure 2E — source data 1 - raw unedited blot.

### GPR183 is required for B-cell trafficking and macrophage-dependent phagocytosis after triple combination treatment

To investigate how the upregulation of GPR183 in target cells could impact their recognition and phagocytosis by M1 macrophages, a single-clone derived Raji-GPR183^KO^ cell was generated by CRISPR/Cas9 gene editing (Figure 3A and source data 1), using previously described procedures (Ribeiro et al., 2021) and co-cultured for 24h with primary M1 macrophages and BMSCs under Nanoshuttle-driven magnetic levitation (Souza et al., 2010) in a conditioned medium to form functional 3D spheroids. Compared to the Raji-GPR183^wt^, the Raji-GPR183^KO^ spheroids harboured a complete blockade of M1 cell infiltration within the multicellular aggregates, both at basal levels and upon treatment with the triplet (Figure 3B). Accordingly, ADCP activity was abrogated in the Raji-GPR183^KO^ cell cultures (Figure 3C) highlighting the critical role of GPR183 in the recruitment of macrophages. ADCC was also compromised, although to a lower extent (Figure 3C). Supporting these results, global inflammatory signature, and especially *CCL20* gene overexpression, were not detected anymore in Raji-GPR183^KO^ co-cultures exposed to triplet therapy (Figure 3D). To understand whether a functional GPR183 was required for the TG-1801+U2 synergistic interaction, Raji cells were exposed for 1h to the GPR183 inhibitor NIBR189 (Gessier et al., 2014), washed out, and co-cultured for 3h with M1-polarized macrophages, in the presence of U2 +/- TG-1801. As shown in Figure 3E, relative ADCP was decreased by 3-fold after GPR183 pharmacological blockade, when compared to untreated Raji cells.

**Figure 3.**
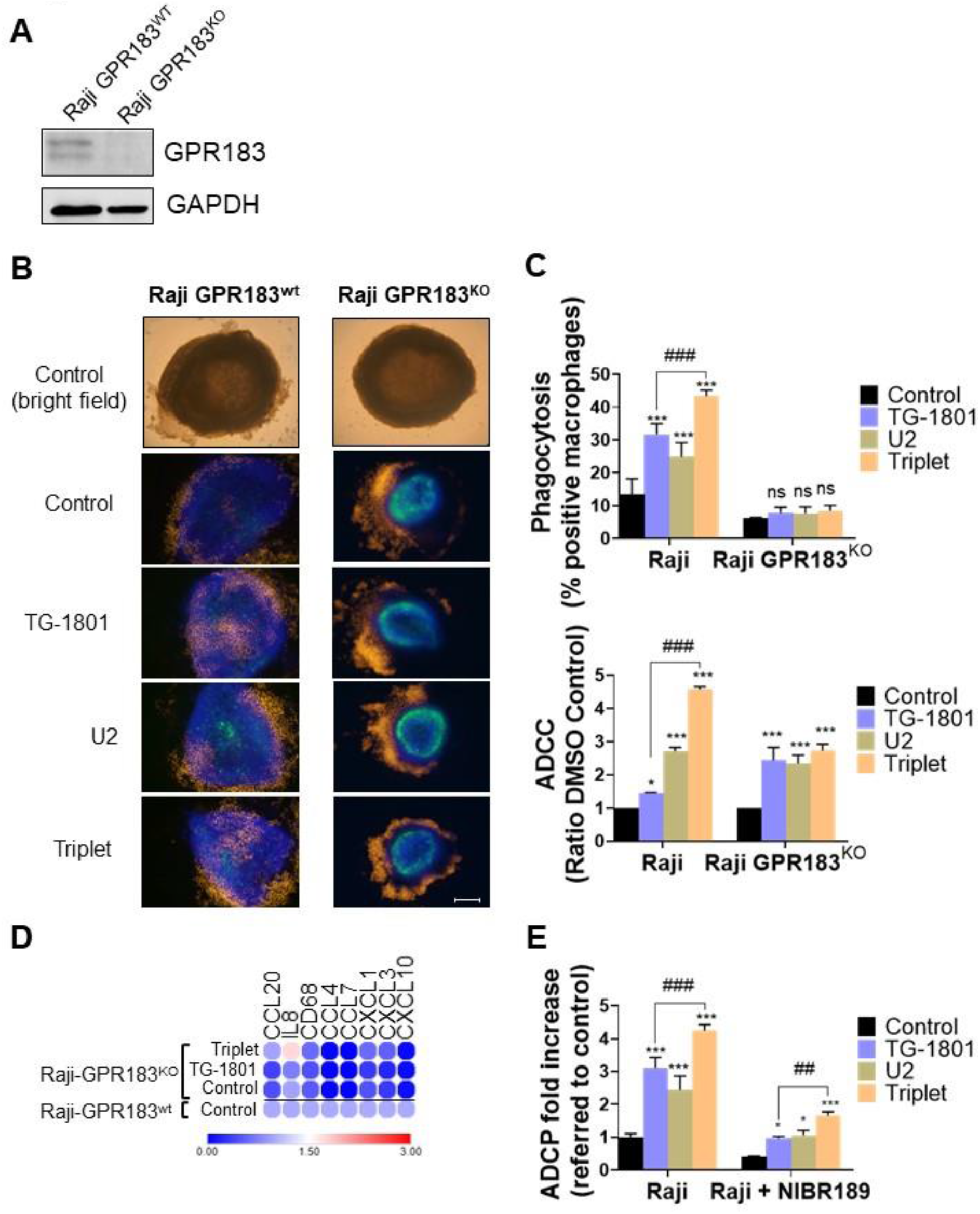
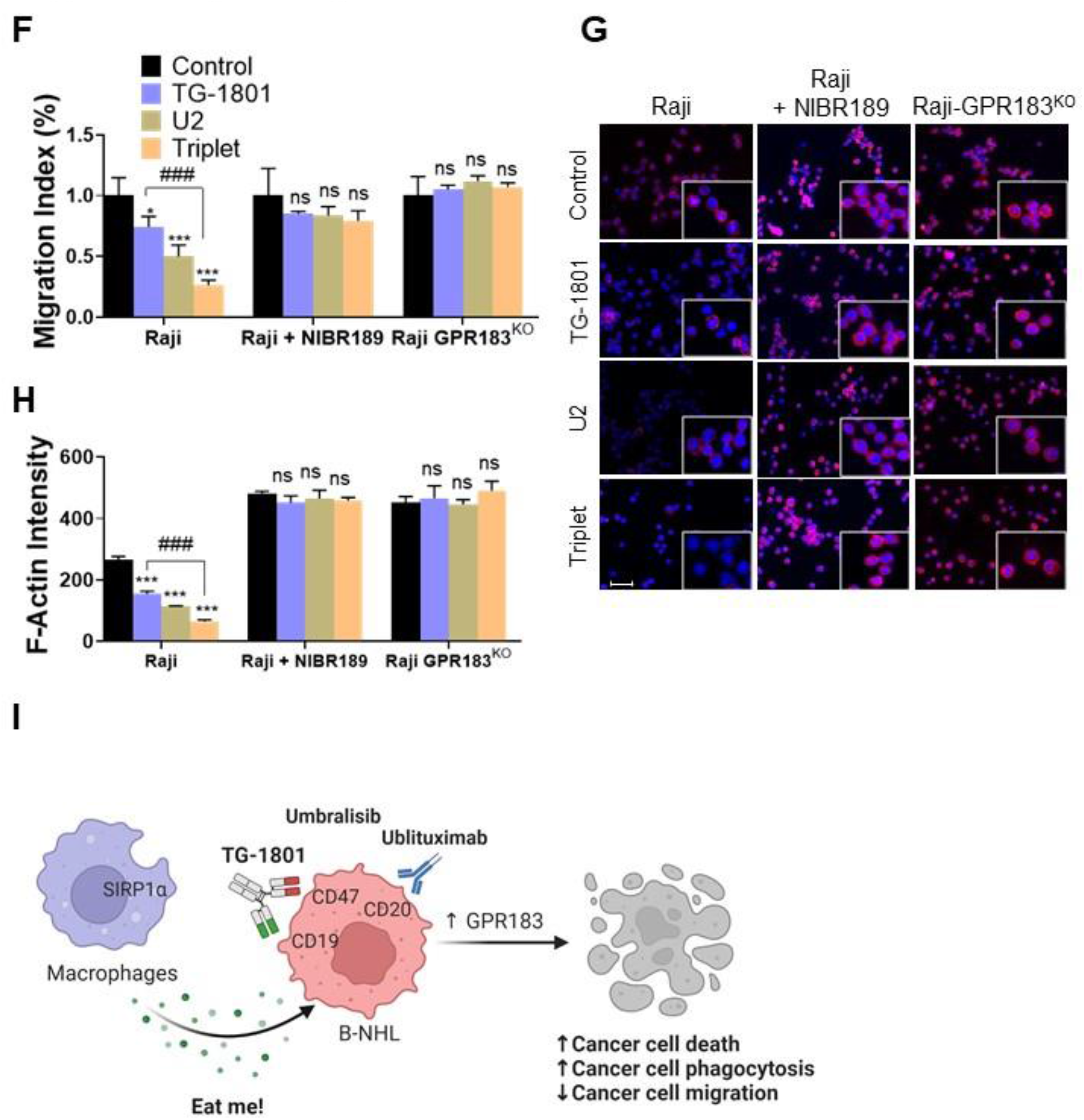
GPR183 activity is required to impair B cell trafficking and to potentiate macrophage-dependent phagocytosis in triplet-treated cells. A) Immunoblot evaluation of GPR183 in both Raji-GPR183^WT^ and Raji-GPR183^KO^ cells. B) Raji-GPR183^WT^ or Raji-GPR183^KO^ 3D spheroid in presence or absence of 10 ng/mL TG-1801 +/- U2 (10 μg/mL ublituximab + 1 μM umbralisib) for one more day. The infiltration of M1 macrophages was evaluated by live-cell red fluorescence (N=3). Scale bar: 500 μm. C) The phagocytosis and cytotoxicity rates were assessed in Raji-GPR183^WT^ and Raji-GPR183^KO^ cultures with pHrodo-stained B cells (top graph) and ADCC (bottom graph), as previously described (N=3). D) Raji-GPR183^WT^ and Raji-GPR183^KO^ were co-cultured with BMSCs, M2-polarized primary macrophages and PBMCs (4:1:1:1) and treated with vehicle, TG-1801 or the triplet combination for 24h. Then, purified CD20+ cells were subjected to RNA extraction and qPCR. Data are presented in fold-change related to the Raji-GPR183^WT^ control (N=3). E) ADCP activities were assessed in Raji cells with or without the GPR183 inhibitor NIBR189 prior to treatment with 10 ng/mL TG-1801 +/- U2 combo (N=3). F) The cell migration index of Raji-GPR183^WT^, Raji-GPR183^KO^ cells exposed to the GPR183 inhibitor NIBR189, in presence or absence of 10 ng/mL TG-1801 +/- U2 combination (N=3). G) F-actin levels were assessed in the different cultures exposed to TG-1801 +/- U2 as in E), followed by staining with a TRITC-labelled phalloidin and direct red fluorescence recording. Nuclei were counterstained with DAPI (blue) (N=3). Scale bar: 50 μm. H) F-actin fluorescence intensity from Raji-GPR183^WT^, Raji-GPR183^KO^, and NIBR189-treated Raji cells in presence or absence of TG-1801 +/- U2 (N=3). Values are expressed as mean ± SD. * *p*<0.05, ** *p*<0.01, *** *p*<0.001, when compared to control group. ^##^ *p*<0.01 and ^##^ *p*<0.001 when compared to TG-1801 alone. ns = non-significant. I) Mechanism of action of the TG-1801/U2 triplet combination therapy in B-NHL. The novel CD47-CD19 bispecific antibody potentiates the anti-tumour activity of the ublituximab-umbralisib regimen through activation of the pro-inflammatory GPR183, thus promoting macrophage-dependent phagocytosis, B-cell cytoskeleton remodelling and B-cell motility. Figure 3A—source data 1 - raw unedited blot.

Since GPR183 is a known antagonist of chemokine-mediated B cell migration (Barroso et al., 2012), a transwell migration assay using recombinant CXCL12 as a chemoattractant was set up, to compare the Raji parental to the Raji-GPR183^KO^ cells and to NIBR189-pretreated cells. Figure 3F shows that cell migration was significantly impaired by both U2 and TG-1801 treatments, and that the combination of the two drugs led to an accentuated inhibition. This effect was completely lost either after GPR183 pharmacological inhibition by NIBR189 or after the genetic deletion of the receptor. Accordingly, F-actin polymerization was decreased by 70% in Raji cells exposed to the triplet treatment and this effect was completely lost in the absence of GPR183 or after NIBR189 treatment (Figures 3G and 3H), in agreement with previous studies that highlighted the relevance of F-actin disruption in target cells in the anti-lymphoma effect of anti-CD47 antibodies (Barbier et al., 2009).

While the synergistic anti-tumour effect of CD47-targeting drugs, when combined with IgG1 antibodies, has classically been related to the inhibition of the “do-not-eat-me” signal, here we propose a novel and additional mechanism based on the overexpression of the proinflammatory receptor GPR183. Our results support a role for GPR183 in the recognition and elimination *in vitro* and *in vivo* of tumour B cells by activated macrophages (Figure 3I). Future studies will be aimed at understanding whether GPR183 could be a biomarker for the activity of therapeutic combinations containing CD47 targeted therapy. Testing whether this discovery can be generalized to other drugs from the same families is underway.

## Materials and Methods

### Cell lines

Five DLBCL (Pfeiffer, TMD-8, HBL-1, SUDHL-5, Karpas-422), three Burkitt lymphoma (Raji, Daudi, Ramos), two Follicular lymphomas (DOHH-2, RL), and one T-cell Acute Lymphoblastic Leukemia (Jurkat) cell lines used in this study were grown in Advanced-RMPI 1640 supplemented with 5% heat-inactivated FBS, 2 mmol/L glutamine, 50 μg/mL penicillin-streptomycin (Thermo Fisher, MA, USA).

### Occupancy assay

Cytofluorimetric quantification of CD47 and CD19 levels in a panel of 10 B-NHL cell lines. Cells were stained with PE-labelled anti-CD47 or anti-CD19 antibodies (Becton Dickinson) and the absolute number of membrane-bound molecules of CD47 or CD19 was estimated using QuantiBRITE PE beads (BD Biosciences) on a FACSCanto II (Becton Dickinson). Data were analysed using FlowJo software package.

For the detection of surface CD47, Raji (CD19+), or Jurkat (CD19-) cells were stained with a phycoerythrin (PE)-labelled anti-CD47 (B6H12 clone) or isotype control antibody (BD Biosciences). The cells were pre-treated for 1 h with TG-1801 or an anti-human CD47 (B6H12 clone) control antibody. For quantification, a total of 10,000 events were acquired on a FACSCanto II (Becton Dickinson). Relative median fluorescence intensity (MFI) was calculated using FlowJo software package as the ratio between CD47 and control signal intensity. Shown are the percentages of occupancy, defined as the decreases in CD47 MFI ratios evoked by anti-CD47-treatment, using untreated cells as a calibrator. B6H12 clone was used as a CD19-independent positive control of CD47 occupancy.

### Peripheral blood mononuclear cells isolation and macrophage polarization

Peripheral blood mononuclear cells (PBMCs) were purified by standard Ficoll-Hypaque (GE Healthcare, UK) gradient centrifugation from buffy coats of human healthy donors and cultured freshly in Advanced-RMPI 1640 supplemented with 5% heat-inactivated FBS, 2 mmol/L glutamine, 50 μg/mL penicillin-streptomycin (Thermo Fisher, MA, USA).

RosetteSep™ Human Monocyte Enrichment Cocktail (Stemcell Technologies, Canada) was used to purify human monocytes from buffy coats following manufacturer specifications. For M1 or M2 macrophage polarization, the selected monocytes were cultured in complete Advanced-RMPI 1640 supplemented with either 20 ng/mL human GM-CSF (PeproTech, RockyHill, NJ), for M1 differentiation, or 20 ng/mL human M-CSF (PeproTech), for M2 differentiation, and incubated for 6 days. On day 6 M0 macrophages were activated with 100 ng/mL human IFN-γ (PeproTech) and 50 ng/mL LPS, for M1 macrophage polarization for 24 h.

### Antibody-dependent cell-mediated cytotoxicity (ADCC) and phagocytosis (ADCP) assays

ADCC activity was assessed in B-cell lymphoma cell lines co-cultured for 4 hours with freshly obtained PBMCs (1:10, target:effector), in the presence of 10 ng/mL TG-1801 +/- U2 dual assets (10 μg/mL ublituximab + 1 μM umbralisib), using and a LDH release assay (Roche). The ADCC was calculated using the following formula:

ADCC percentage = [(sample release – spontaneous release)/(maximal release – spontaneous release)]*100.

Spontaneous release, corresponding to target cells incubated with effector cells without antibody, was defined as 0% cytotoxicity, with maximal release (target cells lysed with 1% Triton X-100) defined as 100% cytotoxicity. The average percentage of ADCC and standard deviations of the triplicates of each experiment were calculated.

ADCP activity was assessed in B-cell lymphoma cell lines co-cultured for 4 hours with M1-polarized macrophages (1:5, target:effector), in the presence of 10 ng/mL TG-1801 +/- U2 dual assets (10 μg/mL ublituximab + 1 μM umbralisib), using and the pHrodo-stained B cells (IncuCyte^®^ pHrodo^®^ Red Cell Labelling Kit for Phagocytosis, Sartorius). Following phagocytosis assay, the non-phagocytosed cells were removed by washing with PBS 2–3 times and phagocytosis was analysed by fluorescent microscopy (EVOS Cell Imaging Systems -Thermo-Fisher)

### Xenograft mouse model and IHC staining

Eight-week-old NOD/SCID IL2Rγ-null (NSG) male and female mice (Janvier Labs, France) were subcutaneously injected with Raji cells and tumour-bearing mice were randomized using GraphPad Prism 9.0 software and assigned to one of the following treatment arms (8-6 mice per group): TG-1801 (5 mg/kg, qw), ublituximab (5 mg/kg, qw), umbralisib (150 mg/kg, bid), ublituximab (5 mg/kg, qw) + umbralisib (150 mg/kg, bid), TG-1801+ublituximab combo, TG-1801+umbralisib combo or the triplet (U2 + TG-1801), or an equal volume of vehicle for 17 days. Tumour volumes were measured each 2-3 days with external callipers. The number of animals used in each of the experimental groups is based on the literature and previous results from the group (Ribeiro et al., 2021). Immunohistochemical staining of representative tumour specimens (N=3) was performed using anti-CD20 (Sigma), anti-GPR183 (Santa Cruz), anti-F4/80 (Abcam), anti-Histone H3-pSer10 (Abcam) and anti-CD56/NCAM-1 (Abcam). Preparations were evaluated using an Olympus microscope and MicroManager software.

IHC signal intensity was quantified in at least 5 pictures of two representative tumour specimens from the Raji xenograft model, using QuPath v.0.2.3 (Queen’s University, Belfast, Northern Ireland). Cell detection was conducted as previously described (Bankhead et al., 2017) using QuPath’s built-in “Positive cell detection” by calculating the per cent of positively stained cells in each field.

### RNA sequencing (RNA-seq) analysis

Two Burkitt lymphoma (BL) cell lines (Daudi and Raji) and two BL primary samples were co-cultured with bone marrow stromal cells (BMSCs), M2-polarized macrophages and PBMCs (4:1:1:1) in the presence of 10 ng/mL TG-1801 +/- U2. After 24h incubation, CD20+ target cells were isolated using the EasySep Human Biotin Positive Selection Kit II (StemCell Technologies, Canada) and the biotinylated anti-CD20 antibody (BioLegend, CA, USA). Purified CD20+ cells, together with representative bulk Raji xenografts with > 95% tumour B cells were subjected to RNA-seq analysis according to previous procedures (Ribeiro et al., 2021). Sequencing data have been deposited at the Gene Expression Omnibus (GEO) of the National Center for Biotechnology Information (GSE199413). Volcano plot showing the most relevant significantly differentially expressed genes between triplet and TG-1801 treatments, with |Log2 fold change| > 1.5 and p-adj value < 0.01 (red dots). Grey, green and blue dots identified genes with insignificant transcriptional and/or statistical variation. Briefly, the raw fastq RNAseq files of each condition were quality checked and gene expression was estimated using Salmon software (https://combine-lab.github.io/salmon/). Differential expression analysis was then carried out using the negative binomial distribution (DESeq2 software, https://bioconductor.org/packages/release/bioc/html/DESeq2.html), accounting for and filtering the effects of the respective controls.

Purified CD20+ cells, together with representative Raji xenografts were subjected to RNA extraction and qPCR validation. Briefly, total RNA was extracted using TRIZOL (Thermo Fisher) following manufacturer’s instructions. One microgram of RNA was retrotranscribed to complementary DNA using moloney murine leukemia virus reverse transcriptase (Thermo Fisher) and random hexamer primers (Roche). mRNA expression was analyzed in duplicate by quantitative real-time PCR and the relative expression of each gene was quantified by the comparative cycle threshold method (ΔΔC_t_) β-actin (Fw: GACGACATGGAGAAAATCTG, Rv: ATGATCTGGGTCATCTTCTC) were used as an endogenous control. The sequences used for the primers are the following GPR183 (Fw: GACTGGAGAATCGGAGATGC, Rv: CAGCAATGAAGCGGTCAATA), CCL20 (Fw: CCAATGAAGGCTGTGACATCA, Rv: AGTCTGTTTTGGATTTGCGCA), IL8 (Fw: AAGGAAAACTGGGTGCAGAG, Rv: GCTTGAAGTTTCACTGGCATC), CD68 (Fw: CCTCCAGCAGAAGGTTGTCT, Rv: CGAAGGGATGCATTCTGAGC), CCL4 (Fw: TTCCTCGCAACTTTGTGGTA, Rv: GCTTGCTTCTTTTGGTTTGG), CCL7 (Fw: TGG AGA GCTACAGAAGGACCA, Rv: GGGTCAGCACAGATCTCCTT), CXCL1 (Fw: CATCCAAAGTGTGAACGTGAA, Rv: CTATGGGGGATGCAGGATT), CXCL3, (Fw: CAAAGTGTGAATGTAAGGTCCCC, Rv: CGGGGTTGAGACAAGCTTTC) and CXCL10 (Fw: CCTGCAAGCCAATTTTGTCCA, Rv: TGGCCTTCGATTCTGGATTCA).

### Generation of Raji-GPR183^KO^ cells

The generation of a CRISPR-Cas9 gene-editing tool was employed to edit the Raji parental cells line to create GPR183 knockout. 0.5 × 10^6^ cells were electroporated on a Nucleofector II device (program A032, Lonza) with 36 pmol SpCas9 Nuclease V3, 44 pmol CRISPR-Cas9 tracRNA ATTO 550, 44 pmol Alt-R CRISPR-Cas9 crRNA <u>Hs.Cas9.GPR183.1.AA</u> (GPR183^KO^ 5’-CAATGAAGCGGTCAATACTC AGG -3’) (IDT-Integrated DNA Technologies). GPR183^KO^ cells were resuspended in 96-well plates with a limiting dilution of 0.3 cells per well. The GPR183^KO^ biallelic clones were confirmed by Sanger Sequencing and western blot. Raji-GPR183^KO^ is available upon request.

### Western blot analysis

Total protein extracts were obtained from cell lines and tumour specimens using RIPA (Sigma-Aldrich) buffer and subjected to SDS-PAGE. Membrane-transferred proteins were revealed by incubating with primary and secondary antibodies followed by chemiluminescence detection using the ECL system (Pierce) and a Fusion FX imaging system (Vilber Lourmat). Band intensity was quantified using Image J software and normalized to housekeeping protein (GAPDH). Values were referred to the indicated control and added below the corresponding band. If not otherwise specified, representative data from N = 2 experiments are shown.

### 3D multicellular spheroid generation

One hundred thousand Raji-GPR183^WT^ or Raji-GPR183^KO^ cells were then stained with Hoechst 33342 blue dye (Invitrogen) and cultivated in a conditional medium with 25,000 StromaNKtert-GFP cells for 2 days to generate the BL 3D spheroids. Then, 25,000 M1-macrophages were stained with PKH26 red-fluorescent dye and added to 3D spheroid in presence or absence of 10 ng/mL TG-1801 +/- U2 (10 μg/mL ublituximab + 1 μM umbralisib) for one more day. The M1-macrophages infiltration was evaluated by live-cell red fluorescence at EVOS Cell Imaging Systems (Thermo-Fisher).

### Transwell migration assay and F-actin staining

Briefly, Raji-GPR183^WT^, Raji-GPR183^KO^ and Raji parental cells exposed to the GPR183 inhibitor NIBR189 (Sigma-Aldrich, Germany) were cultured for 1 h in culture medium not containing foetal bovine serum but supplemented with 0.5% bovine serum albumin (Sigma-Aldrich), in the presence or absence of 10 ng/mL TG-1801 +/- U2 combination, and analysed for CXCL12-dependent chemotaxis, as previously described (Balsas et al., 2017). Values were referred to cells cultured without CXCL12. F-actin levels were assessed after exposure to TG-1801 +/- U2, followed by staining with a TRITC-labelled phalloidin and direct red fluorescence recording.

### Ethical issues

Animals were handled following protocols approved by the Animal Ethics Committee of the University of Barcelona (registry num. 38/18).

Institutional Review Board approvals for the study protocol (ref PI-20-040), amendments, and written informed consent documents from BL patients and healthy donors were obtained prior to study initiation. Study procedures were conducted in accordance with the Declaration of Helsinki. Buffy coats were provided by the Blood and Tissue Bank of Catalonia (agreement NE-A1-IJC).

### Statistical analysis

Presented data are the mean ± SD or SEM of 3 independent experiments. All statistical analyses were done by using GraphPad Prism 9.0 software (GraphPad Software). Comparison between 2 groups of samples was evaluated by nonparametric Mann–Whitney test to determine how the response is affected by 2 factors. Pearson test was used to assess the statistical significance of correlation. Results were considered statistically significant when *p*-value < 0.05.

## Acknowledgments

We are very grateful to Salvador Sánchez Vinces, at Sao Francisco University/Brazil for his precious help on bioinformatic. This study was financially supported by TG Therapeutics, Fondo de Investigación Sanitaria PI18/01383, European Regional Development Fund (ERDF) “Una manera de hacer Europa” (to GR). JCS and MFS were recipients of a Sara Borrell research contract (CD19/00228) and a predoctoral fellowship (FI19/00338) from Instituto de Salud Carlos III, respectively. MA was a fellow of PROTEOblood, a project co-financed by the European Regional Development Fund (ERDF) through the Interreg V-A Spain-France-Andorra (POCTEFA) program (EFA360/19). This work was carried out under the CERCA Program (Generalitat de Catalunya).

---

**Figure 1- figure supplement legend.**
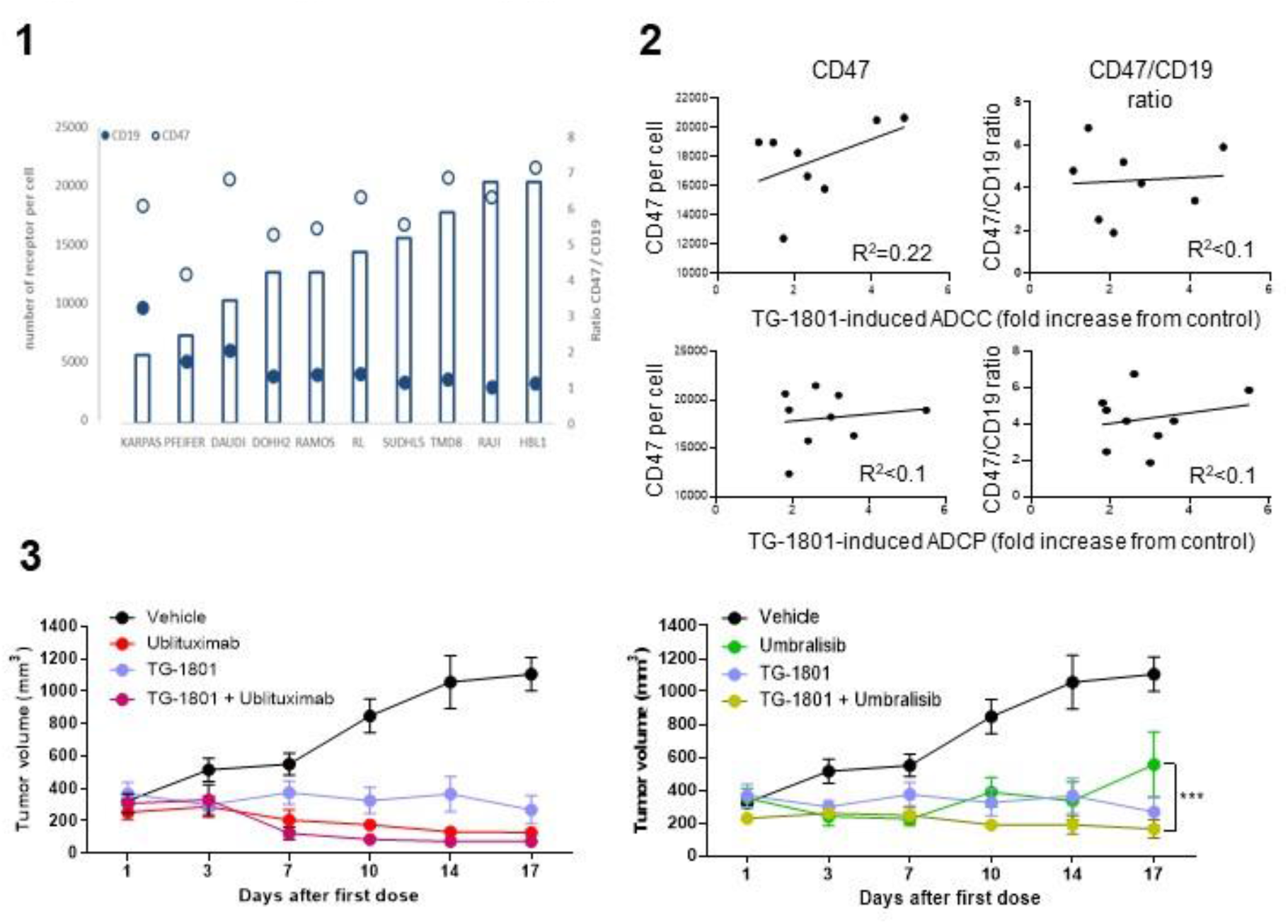
1) Cytofluorimetric quantification of CD47 and CD19 levels in a panel of 10 B-NH cell lines. 2) CD47 and CD19 expression levels, as well as their ratios, were plotted against the corresponding phagocytosis and ADCC quantification for each cell line described in 1), and the correlation coefficient was calculated using GraphPad Prism software. 3) NSG mice were subcutaneously injected with Raji cells and tumour-bearing animals were randomly assigned to one of the following treatment arms (8-6 mice per group): TG-1801 (5 mg/kg, qw), ublituximab (5 mg/kg, qw), umbralisib (150 mg/kg, bid), TG-1801+ublituximab combo (left panel) or TG-1801+umbralisib combo (right panel), or an equal volume of vehicle, for 17 days. Tumour volumes were recorded each 2-3 days using external callipers. *** *p*<0.001.

